# The tetraspanin TSPAN5 regulates AMPARs exocytosis by interacting with the AP-4 complex

**DOI:** 10.1101/2022.01.04.474946

**Authors:** Moretto Edoardo, Longatti Anna, Federico Miozzo, Caroline Bonnet, Francoise Coussen, Fanny Jaudon, Lorenzo A. Cingolani, Passafaro Maria

**Author notes:** These authors contributed equally to this work (Moretto Edoardo, Longatti Anna). **Impact statement** In mature neurons, TSPAN5 assembles a complex with AP-4 and Stargazin to promote the exocytosis of newly synthesised GluA2-containing AMPA receptors.

## Abstract

Intracellular trafficking of AMPA receptors is a tightly regulated process which involves several adaptor proteins, and is crucial for the activity of excitatory synapses in both basal conditions and during synaptic plasticity. We found that, in rat hippocampal neurons, an intracellular pool of the tetraspanin TSPAN5 specifically promotes exocytosis of newly synthesised GluA2-containing AMPA receptors without affecting their internalisation. TSPAN5 mediates this function by interacting with AP-4 and Stargazin and possibly using recycling endosomes as a delivery route. This work highlights TSPAN5 as a new adaptor regulating AMPA receptor trafficking. In addition, it provides a possible mechanism for the intellectual disability symptoms that occur in AP-4 deficiency syndrome.

## Introduction

Tetraspanins are transmembrane proteins conserved in metazoans that present four transmembrane domains (TM), a small and a large extracellular loop (SEL and LEL, respectively), and intracellular N- and C-termini (Berditchevski, 2001). Tetraspanins have the peculiar ability to organise Tetraspanin Enriched Microdomains (TEMs), membrane domains in which they accumulate (Charrin et al., 2002). Tetraspanins have been proposed to function as molecular facilitators by promoting physical proximity between proteins that belong to signalling complexes (Charrin et al., 2014; Maecker et al., 1997). To date, 33 tetraspanins have been described in mammals, with functions in cell-cell adhesion, sperm-egg fusion, cell motility and proliferation (Hemler, 2005). TSPAN5 is part of the C8 subgroup of tetraspanins and was previously shown to regulate the intracellular trafficking and activity of the protease ADAM-10 (Dornier et al., 2012a; Eschenbrenner et al., 2020; Haining et al., 2012a; Jouannet et al., 2016; Noy et al., 2016a; Saint-Pol et al., 2017).

A previous study from our laboratory showed that, in hippocampal pyramidal neurons, TSPAN5 is enriched in dendritic spines and promotes their morphological maturation during synaptogenesis (Moretto et al., 2019a). This action is mediated by controlling the surface mobility of the postsynaptic adhesion molecule neuroligin-1 via an interaction occurring on the plasma membrane. A few other studies have investigated the function of tetraspanins at the synapse identifying their role in intracellular trafficking of neurotransmitter receptors in neurons (Bassani et al., 2012; Lee et al., 2017; Murru et al., 2017, 2018).

Here, we report a significant increase in the intracellular pool of TSPAN5 in dendritic spines upon neuronal maturation. We demonstrate that this pool does not participate in dendritic spine maturation but has the main function to control surface delivery of GluA2-containing α-amino-3-hydroxy-5-methyl-4-isoxazolepropionic acid (AMPA) receptors (AMPARs). AMPARs are tetrameric complexes that mediate most of the fast excitatory transmission in response to the neurotransmitter glutamate in neurons (Henley and Wilkinson, 2016). AMPAR surface levels are directly responsible for synapse weakening or strengthening during synaptic plasticity (Huganir and Nicoll, 2013); their intracellular trafficking is an extremely complex phenomenon which involves several auxiliary proteins (Anggono and Huganir, 2012; Moretto and Passafaro, 2018) that can be impaired in neurological and neurodevelopmental disorders (Henley and Wilkinson, 2016; Moretto et al., 2016, 2017).

Importantly, we found that TSPAN5 exerts this function by interacting with AP-4, a member of the adaptor protein complex family (Boehm and Bonifacino, 2001a; Bonifacino, 2014a; Robinson and Bonifacino, 2001a), which coding genes are mutated in a syndrome characterised by spastic paraplegia and intellectual disability (Sanger et al., 2019a).

Our data identify a novel function of TSPAN5 at the synapse and highlight a possible mechanism for intellectual disability symptoms in the AP-4 deficiency syndrome.

## Results

### TSPAN5 intracellular pool interacts with AP-4 complex

In our previous work (Moretto et al., 2019a), we observed the existence of a substantial intracellular pool of TSPAN5 in mature neurons. We thus performed crosslinking experiments using bis(sulfosuccinimidyl)suberate (BS3) on rat cultured hippocampal neurons. This crosslinker is not permeable to membranes and, as such, if applied to living cells will only crosslink plasma membrane proteins which will appear as high molecular weight bands upon western blot analysis. In contrast, the intracellular pool will not be crosslinked running at the expected molecular weight. This allows one to distinguish between the intracellular and plasma membrane pools. We analysed TSPAN5 levels in these experiments and looked at two different time points: DIV12 and DIV19. At DIV12 synaptogenesis is prominent in rat cultured neurons (Chanda et al., 2017) while at DIV19, primary neurons are considered functionally mature. We observed an increase in the intracellular levels of TSPAN5 from DIV12 to DIV19, which was not followed by a concomitant increase in plasma membrane levels (Fig 1A), suggesting that increased intracellular levels of TSPAN5 does not necessarily imply increased delivery of this protein to the plasma membrane. The transferrin receptor showed a more stable distribution across these time points. To test if the increase of intracellular TSPAN5 could be related to a different function compared to its previously described role in dendritic spines maturation (Moretto et al., 2019a), we transfected cultured hippocampal neurons with Scrambled, Sh-TSPAN5 and Rescue constructs at DIV13. TSPAN5 reduction at this time point would unlikely affect dendritic spine maturation as synaptogenesis is already underway. We analysed dendritic spine density and morphology at DIV21 (Fig 1B) and observed that dendritic spine density was reduced, but to a lower extent compared to our previous observations when knocking down TSPAN5 at DIV5 (20% reduction compared to more than 65%, respectively) (Moretto et al., 2019a). Even more interestingly, the morphology of dendritic spines was completely unaffected by TSPAN5 knockdown at this time point. In contrast, our previous results had shown a strong reduction (50%) in mature mushroom dendritic spines in favour of less mature thin dendritic spines when TSPAN5 levels are downregulated from DIV5 (Moretto et al., 2019a). These data support a more prominent role of TSPAN5 for dendritic spine maturation at early stages of development and suggests that TSPAN5 might be involved in other functions at more mature stages. We decided to explore whether the intracellular TSPAN5 could have a role in regulating intracellular trafficking given previous evidence on the role of this and other tetraspanins (Bassani et al., 2012; Dornier et al., 2012b; Haining et al., 2012b; Jouannet et al., 2016; Noy et al., 2016b; Saint-Pol et al., 2017).

**Figure 1.**
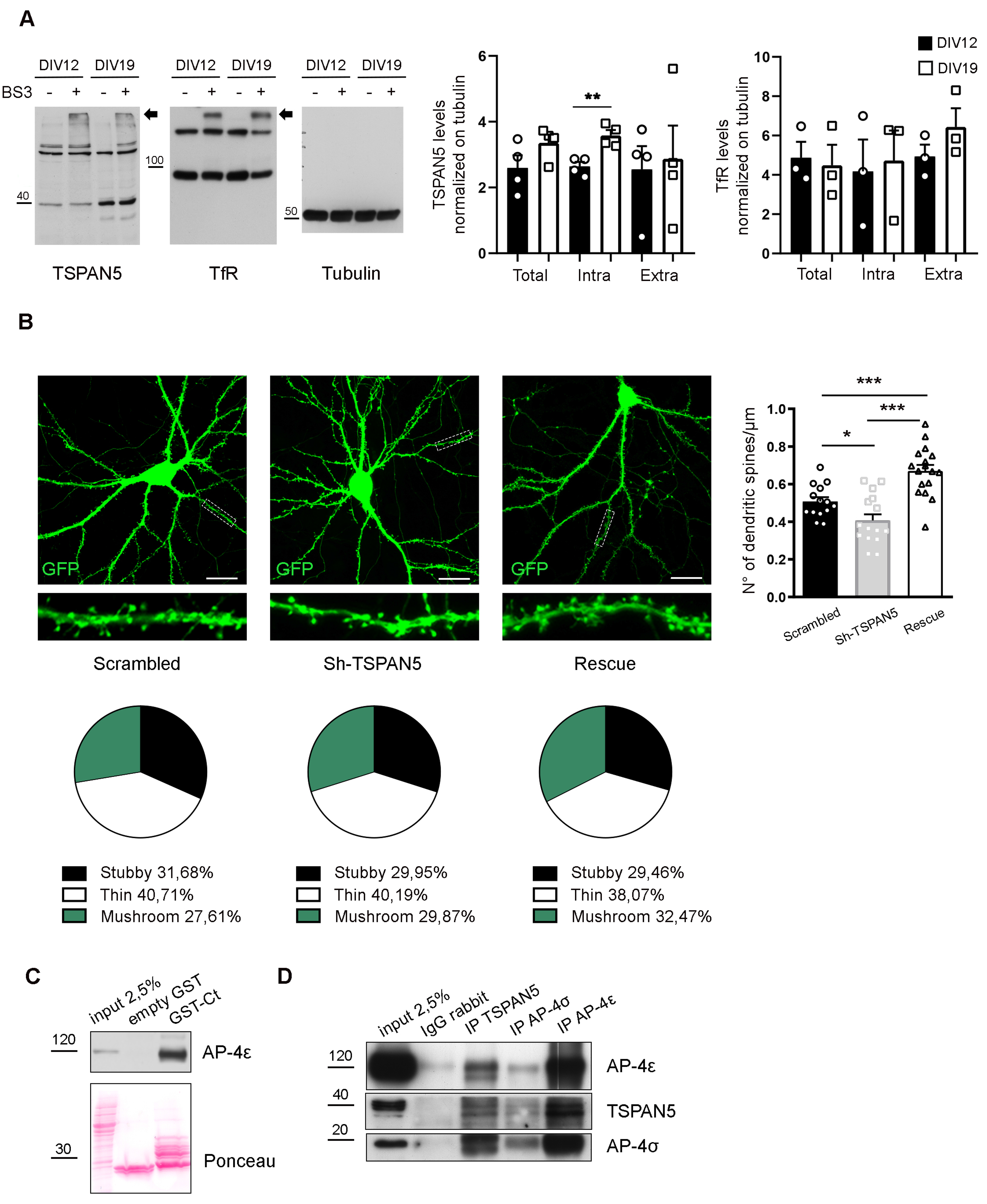
TSPAN5 intracellular pool interacts with AP-4 complex. **(A)** BS3 crosslinking experiment on cultured rat hippocampal neurons at DIV 12 and 19 blotted for TSPAN5, Transferrin receptor (TfR) and Tubulin. Arrows indicate the higher molecular weight bands present in the BS3 + lanes that represent the plasma membrane pool of the proteins. Tubulin was used as loading control; Transferrin Receptor (TfR) was used as crosslinking positive control. (TSPAN5: Total/Tubulin: DIV12 2.599±0.38, DIV19 3.357±0.25; Intra/Tubulin: DIV12 2.643±0.14, DIV19 3.582±0.16; Extra/Tubulin: DIV12 2.552±0.70, DIV19 2.871±1.01; TfR: Total/Tubulin: DIV12 4.87±0.81, DIV19 4.48±1.05; Intra/Tubulin: DIV12 4.18±1.61, DIV19 4.73±1.53; Extra/Tubulin: DIV12 4.94±0.6, DIV19 6.44±0.95). n = 3/4 independent cultures per condition. **(B)** Left panels: confocal images of DIV20 cultured rat hippocampal neurons transfected at DIV12 with either Scrambled, Sh-TSPAN5 or Rescue constructs, all co-expressing GFP. Inserts show higher magnification of the dendrites highlighted in white. Scale bar = 20 μm. Right panel: quantification of dendritic spine density represented as histograms. Dendritic spine density (N° of dendritic spines/μm: Scrambled 0.51±0.02; Sh-TSPAN5 0.41±0.03; Rescue 0.67±0.03). Pie charts (bottom panels) show quantification of dendritic spine morphology. Dendritic spine morphology (%: Stubby: Scrambled 31.20±1.52, Sh-TSPAN5 30.25±2.02, Rescue 29.46±1.38; Thin: Scrambled 40.10±2.45, Sh-TSPAN5 40.51±1.96, Rescue 38.07±2.5; Mushroom: Scrambled 27.15±2.25, Sh-TSPAN5 29.24, Rescue 32.49±2.44). n = Scrambled, 14; Sh-TSPAN5, 16; Rescue, 17 neurons. **(C)** GST-pulldown experiment on adult rat hippocampus and cortex lysates using empty GST or GST fused to TSPAN5 C-terminus (GST-Ct). Input: 2.5% of pulldown volume. Blots probed for AP-4ε. Red Ponceau shows the GST-bound fragments. **(D)** Co-immunoprecipitation experiment on adult rat hippocampus and cortex lysates. Input: 2.5% of the immunoprecipitated volume. Immunoprecipitation: α-rabbit IgG, α-TSPAN5, α-AP4σ or α-AP4ε. Blots probed for TSPAN5, AP-4σ and AP-4ε. Values represent the mean ± SEM. * = p<0.05, ** = p<0.01, *** = p<0.001.

Given that TSPAN5 in the intracellular pool would have its SEL and LEL facing the lumen of vesicles, we hypothesised that this fraction of TSPAN5 could participate in intracellular trafficking through interactions of its cytosol-exposed C-terminus as previously shown for other tetraspanins (Bassani et al., 2012). We thus decided to perform a yeast two-hybrid screen using the C-terminal tail of TSPAN5 as bait. Among the clones identified (the full list is presented in the supplementary figure related to Figure 1), four of them coded for amminoacids 1-102 of the protein AP-4σ. AP-4σ is one of the subunits of the adaptor protein complex AP-4 (Boehm and Bonifacino, 2001b; Bonifacino, 2014a; Robinson and Bonifacino, 2001b). This complex is an obligate tetramer of four different subunits (β, μ, ε and σ) which readily assemble and are almost undetectable as single subunits (Hirst et al., 2013). The AP-4 complex has been previously shown to participate in intracellular trafficking of transmembrane proteins including AMPARs via direct interaction with the auxiliary AMPAR subunit Stargazin (Matsuda et al., 2008). We validated the interaction between TSPAN5 and AP-4 by GST pulldown on rat brain lysates (cortices and hippocampi) using the C-terminus of TSPAN5 fused to GST (GST-Ct) which precipitated AP-4ε (Fig 1C), one of the subunits of the AP-4 complex. In addition, we confirmed the interaction via co-immunoprecipitation experiments by immunoprecipitating TSPAN5, AP-4σ or AP-4ε from rat brain lysates (cortices and hippocampi) and found that all three proteins were associated (Fig 1D).

### TSPAN5 can form a complex with GluA2 and Stargazin and localises in recycling endosomes

Given the previously shown interaction of AP-4 with Stargazin and AMPARs (Matsuda et al., 2013a), we performed GST-pulldown experiments to investigate whether TSPAN5 could be part of the same protein complex. By using the C-terminus of TSPAN5 for GST-pulldown experiments on rat brain lysates, we were able to confirm that the C-terminal tail of TSPAN5 is sufficient to precipitate Stargazin, GluA1 and GluA2/3 (Fig 2A). The NMDA receptor subunit 2A instead was not detected in the precipitates, supporting the specificity of the interaction. To further characterize the interaction, we performed GST pulldown using the C-terminal tail of Stargazin, which was previously identified to be the region responsible for the interaction with AP-4 (Matsuda et al., 2008). With this experiment, we detected GluA2/3 and TSPAN5 in the precipitate (Fig 2B). As negative control, CD81, another member of the tetraspanin family was not precipitated by the GST-Ct-Stargazin.

**Figure 2.**
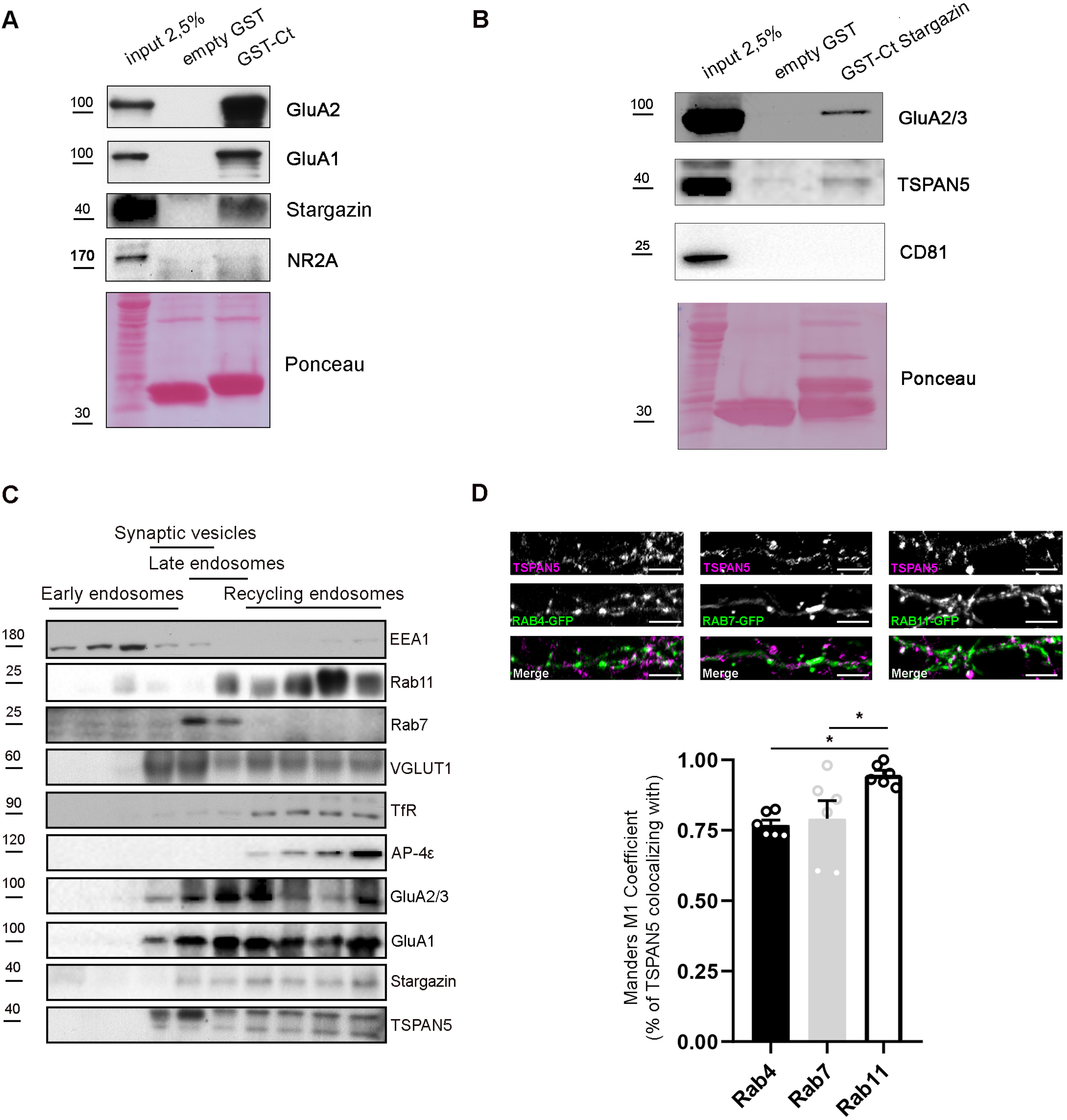
TSPAN5 forms a complex with GluA2 and Stargazin and localises in recycling endosomes. **(A)** GST-pulldown experiments on adult rat hippocampus and cortex lysates using empty-GST or GST fused to the C-terminus of TSPAN5 (GST-Ct). Input: 2.5% of pulldown volume. Blots probed for GluA2/3, GluA1, Stargazin and NMDAR subunit GluN2A. **(B)** GST-pulldown experiments on adult rat hippocampus and cortex lysates using empty-GST or GST-fused to the C-terminal of Stargazin (GST-Ct Stargazin). Input: 2.5% of pulldown volume. Blots probed for GluA2/3, TSPAN5 and CD81. **(C)** Vesicles fractionation from synaptosomes obtained from adult rat hippocampus and cortex. Ten isovolumetric fractions were isolated. Blots were probed for: EEA1 for early endosomes, Rab7 for late endosomes, VGLUT1 for synaptic vesicles, RAB11 and TfR for recycling endosomes, AP-4ε, GluA1, GluA2/3, Stargazin and TSPAN5. **(D)** Top panel: confocal images of DIV20 cultured rat hippocampal neurons transfected at DIV12 with plasmids encoding either Rab4-GFP, Rab7-GFP or Rab11-GFP and immunolabeled for TSPAN5 (magenta). Bottom panel: quantification of TSPAN5 co-localisation (Mander’s M1 coefficient) with RAB4-GFP, RAB7-GFP and RAB11-GFP. (Mander’s M1 coefficient: Rab4, 0.77±0.02; Rab7, 0.79±0.06; 0.94±0.02). n = 6 neurons per condition. Scale bar = 5 μm. Values represent the mean ± SEM. * = p<0.05, ** = p<0.01, *** = p<0.001.

We next addressed where this association was taking place. Given that the intracellular pool of TSPAN5 must reside in intracellular vesicles, we prepared synaptosomes from rat brains (cortices and hippocampi) and loaded their content on a linear sucrose gradient to separate different populations of vesicles (Rao et al., 2011) (Fig 2C). We observed a significant enrichment of AP-4ε, TSPAN5, Stargazin, GluA1 and GluA2/3 in the heaviest fractions which are positive for recycling endosomes markers Rab11 and the Transferrin receptor. This experiment suggests that the intracellular pool of TSPAN5 could associate with AP-4, Stargazin, GluA1 and GluA2 in recycling endosomes. We also analysed the localisation of TSPAN5 in cultured hippocampal neurons by evaluating its colocalisation with overexpressed GFP-tagged Rabs: Rab4, Rab7 and Rab11, markers of early, late and recycling endosomes respectively (Fig 2D). As expected for a transmembrane protein that also localises in the plasma membrane, TSPAN5 had a high level of colocalisation with all three Rabs analysed; however, colocalisation with Rab11 positive endosomes was significantly higher than with the other two Rabs. Taken together, these findings suggest that TSPAN5, AP-4, Stargazin and AMPARs could be associated in Rab11 positive organelles.

### TSPAN5 depletion affects surface and total levels of AMPAR subunits GluA2 and GluA1

Our data so far suggested that TSPAN5 could participate in the intracellular trafficking of AMPARs, a tightly regulated process that is crucial to maintain a correct level of receptors at the synapse membrane and ensure efficient synaptic transmission (Anggono and Huganir, 2012; Hanley, 2010).

To investigate this possibility, we transfected rat hippocampal neurons at DIV12 with either Scrambled, Sh-TSPAN5 or Rescue constructs and measured the surface levels of the two most abundant subunits of AMPARs GluA2 (Fig 3A) and GluA1 (Fig 3B) at DIV20. We observed that knockdown of TSPAN5 induced a reduction of surface GluA2 levels that was reversed in the Rescue condition (Fig 3A). In contrast, GluA1 appeared to be increased upon TSPAN5 knockdown (Fig 3B), an effect that was reversed in the Rescue condition. We also analysed surface levels of both GluA2 and GluA1 specifically in the postsynaptic compartment, by restricting the analysis on dendritic spines or dendritic shafts as identified in the GFP channel by morphological criteria. The reduction of GluA2 and the increase in GluA1 were present in both compartments (Fig 3A), suggesting that this effect is not restricted to the synapse.

**Figure 3.**
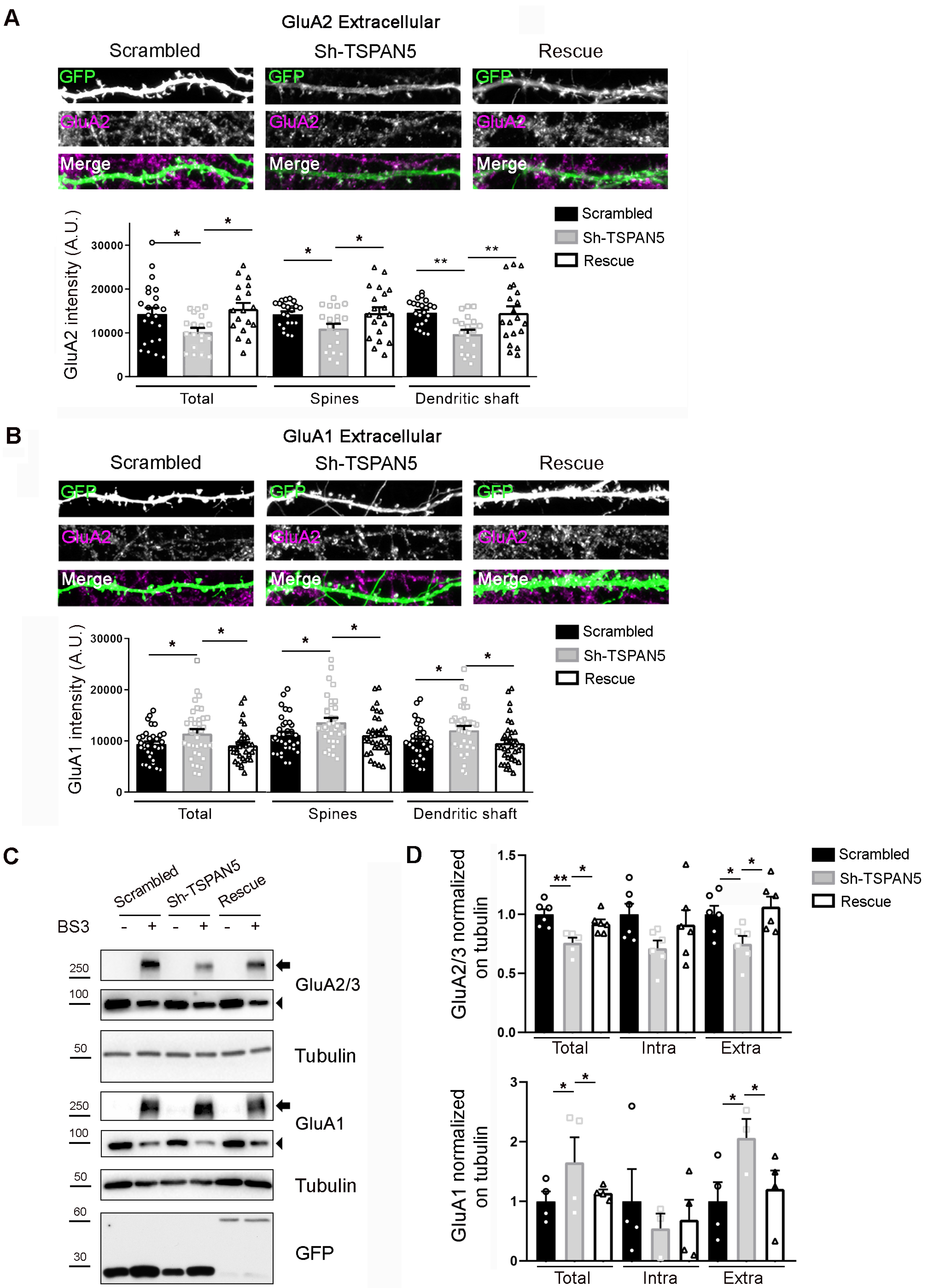
TSPAN5 depletion affects surface and total levels of AMPAR subunits GluA2 and GluA1. **(A)** Top panel: Confocal images of dendrites from cultured rat hippocampal neurons transfected at DIV12 with either Scrambled, Sh-TSPAN5 or Rescue constructs, all co-expressing GFP immunostained at DIV20 with an antibody against an extracellular epitope of GluA2 (magenta) in non-permeabilising condition. Boxes are 20 μm wide. Full neurons are shown in the Supplementary figure relative to figure 3. Bottom panel: quantification of surface GluA2 signal mean intensity on the whole GFP positive area (GluA2 intensity (A.U.) Total: Scrambled 14302±1430, Sh-TSPAN5 10250±884, Rescue 15476±1352); GluA2 mean intensity in dendritic spines (Scrambled 14290±593; Sh-TSPAN5 11006±1055; Rescue 14544±1293); GluA2 mean intensity in dendritic shafts (Scrambled 14579±610; Sh-TSPAN5 9730±921; Rescue 14512±1482). n = Scrambled, 23; Sh-TSPAN5, 19; Rescue, 18 neurons. **(B)** top panel: Confocal images of dendrites from cultured rat hippocampal neurons transfected at DIV12 with either Scrambled, Sh-TSPAN or Rescue constructs, all co-expressing GFP immunostained at DIV20 with an antibody against an extracellular epitope of GluA1 (magenta) in non-permeabilising condition. Boxes are 20 μm wide. Full neurons are shown in the Supplementary figure relative to figure 3. Bottom panel: quantification of GluA1 signal mean intensity on the whole GFP positive area (GluA1 intensity (A.U.) total: Scrambled 9404±494, Sh-TSPAN5 11492±817, Rescue 9167±565); GluA1 mean intensity in dendritic spines (Scrambled 11232±599; Sh-TSPAN5 13711±831; Rescue 11185±634); GluA1 mean intensity in dendritic shafts (Scrambled 9914±563; Sh-TSPAN5 12128±808; Rescue 9610±678). n = Scrambled, 35; Sh-TSPAN5, 36; Rescue, 35 neurons. **(C)** BS3 crosslinking on DIV20 cultured rat hippocampal neurons infected at DIV12 with lentiviral particles encoding for Scrambled, Sh-TSPAN5 or Rescue all co-expressing GFP. Blots probed for AMPARs subunits GluA2/3 and GluA1. Tubulin was used as a loading control, GFP was used as a control of infection. Arrowheads indicate total and intracellular bands, arrows indicate crosslinked plasma membrane bands. Full blots are shown in the Supplementary figure relative to figure 3. **(D)** Quantification relative to panel C. (GluA2/3: Total/Tubulin: Scrambled 1±0.04, Sh-TSPAN5 0.76±0.06, Rescue 0.91±0.04; Intra/Tubulin: Scrambled 1±0.09, Sh-TSPAN5 0.71±0.09, Rescue 0.91±0.14; Extra/Tubulin: Scrambled 1±0.06, Sh-TSPAN5 0.75±0.09, Rescue 1.08±0.08; GluA1: Total/Tubulin: Scrambled 1±0.15, Sh-TSPAN5 1.65±0.22, Rescue 1.14±0.0; Intra/Tubulin: Scrambled 1±0.48, Sh-TSPAN5 0.5±0.4, Rescue 0.69±0.44; Extra/Tubulin: Scrambled 1±0.29, Sh-TSPAN5 2.06±0.14, Rescue 1.21±0.23). n = 4/6 independent cultures. Values represent the mean ± SEM. * = p<0.05, ** = p<0.01, *** = p<0.001.

We confirmed these results by using BS3 crosslinking in hippocampal neurons that were transduced with lentiviral particles carrying Scrambled, Sh-TSPAN5 or Rescue DNA (Fig 3 C). In these experiments, we observed a significant reduction in the plasma membrane and in the total levels of GluA2/3, possibly suggesting an increased degradation of the receptor in addition to its reduced plasma membrane localisation. Similarly, the increase in GluA1 was observed both in the plasma membrane fraction and in the total level suggesting a potential compensatory effect driven by increased protein synthesis or reduced degradation.

### TSPAN5 and AP-4 regulate surface GluA2 levels without affecting its internalisation

To investigate whether the TSPAN5-AP4 complex is responsible for the regulation of GluA2 surface levels, we designed guide RNAs (gRNAs) to knockdown AP-4 β and ε via CRISPR/Cas9 and tested them by transfection in N2A cells followed by Real-Time PCR (Fig 4A). As previously described (Matsuda et al., 2008; Hirst et al., 2013), reduction of the levels of one subunit similarly affected the other subunits. Lentiviral particles coding for these CRISPR/Cas9 knockdown constructs showed efficient reduction in the levels of AP4 β and ε in cultured rat hippocampal neurons, as assessed by RT-PCR (Fig 4B). We then transduced cultured rat hippocampal neurons at DIV12 with lentiviral particles coding for either a control, AP-4 β or AP-4 ε gRNAs and simultaneously with lentiviral particles coding for Scrambled or TSPAN5 shRNA. We subsequently analysed the surface levels of GluA2 at DIV20 and observed that the knockdown of either AP-4 or TSPAN5 reduced such levels to the same extent and that the simultaneous knockdown of TSPAN5 and AP-4 did not induce any further reduction (Fig 4C), supporting the hypothesis that the two proteins participate in the same pathway.

**Figure 4.**
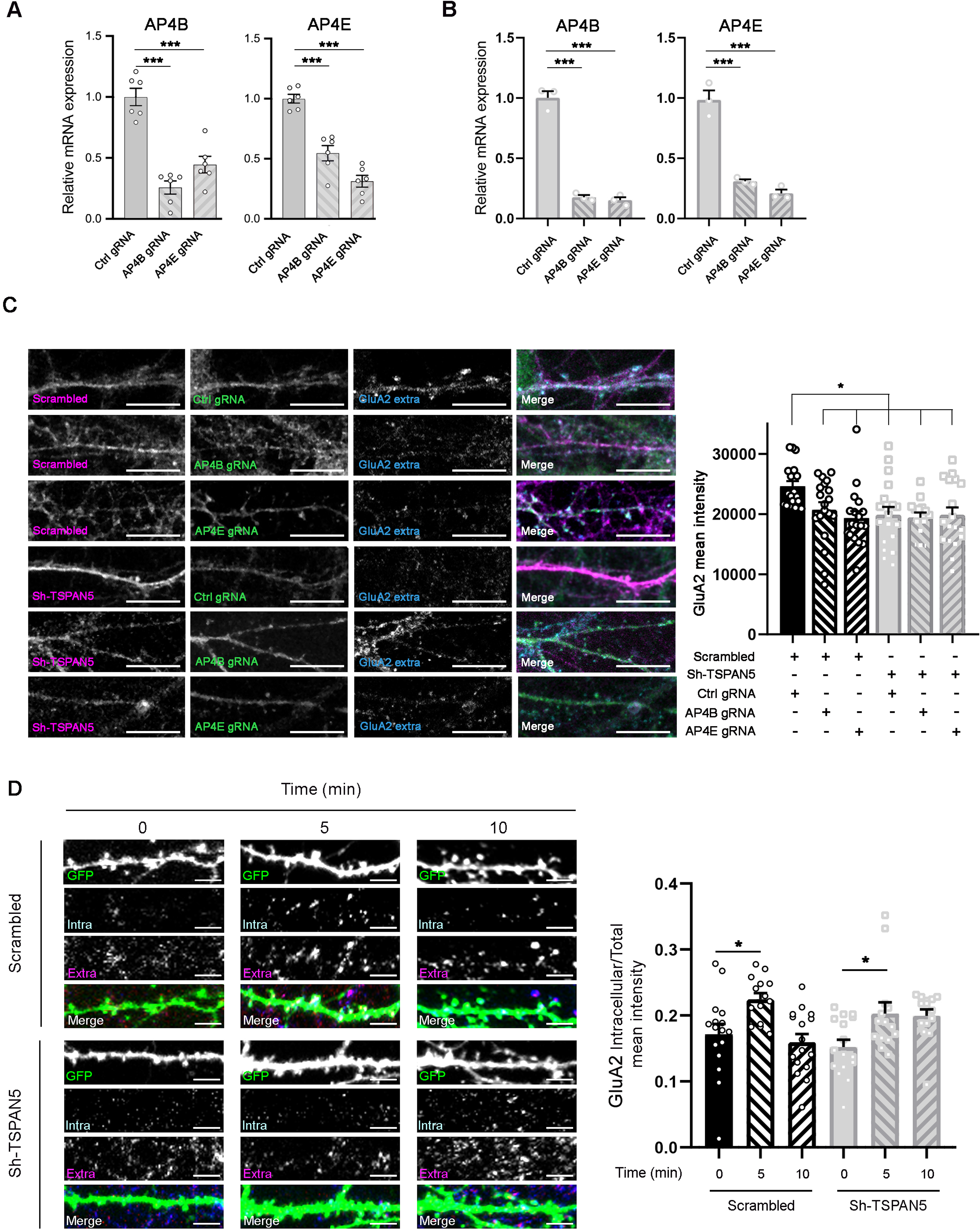
TSPAN5 and AP-4 regulate surface GluA2 levels without affecting its internalisation. **(A)** RealTime PCR quantification of the relative AP4B (left panel) and AP4E (right panel) mRNA levels in N2A cells transfected with plasmids coding for CRISPR/Cas9 and either a control guide RNA or guide RNAs directed against AP4B or AP4E, respectively. mRNA levels normalised to levels in Ctrl gRNA: AP4B (Ctrl gRNA 1±0.07; AP4B gRNA 0.26±0.05; AP4E gRNA 0.45±0.07); AP4E (Ctrl gRNA 1±0.04; AP4B gRNA 0.55±0.06; AP4E gRNA 0.31±0.03). n = 6 from 3 independent experiments. **(B)** RealTime PCR quantification of the relative AP4B (left panel) and AP4E (right panel) mRNA levels in DIV20 cultured rat hippocampal neurons infected at DIV12 with lentiviral particles coding for CRISPR/Cas9 and either a control guide RNA or guide RNAs directed against AP4B or AP4E, respectively. mRNA levels normalised to levels in Ctrl gRNA: AP4B (Ctrl gRNA 1±0.05; AP4B gRNA 0.18±0.02; AP4E gRNA 0.15±0.02); AP4E (Ctrl gRNA 1±0.08; AP4B gRNA 0.31±0.02; AP4E gRNA 0.21±0.03). n = 3 independent experiments. **(C)** Left panel: Confocal images of DIV20 culture rat hippocampal neurons infected at DIV12 with lentiviral particles coding for an mCherry (magenta) and either Scrambled or Sh-TSPAN5 and with lentiviral particles coding for a GFP (green) and either a control guide RNA (Ctrl gRNA) or gRNAs directed against AP4B (AP4B gRNA) or AP4E (AP4E gRNA), respectively, and immunostained at DIV20 with an antibody against an extracellular epitope of GluA2 (cyan). Right panel: Quantification of the intensity of the GluA2 signal: GluA2 mean intensity (Scrambled-Ctrl gRNA 24627±840; Scrambled-AP4B gRNA 20737±1236; Scrambled-AP4E gRNA 19339±1165; Sh-TSPAN5-Ctrl gRNA 19864±1331; Sh-TSPAN5-AP4B gRNA 19407±836; Sh-TSPAN5-AP4E gRNA 19836±1279). n = 18 neurons from 3 independent experiments. Scale bar = 10 μm. **(D)** Left panel: confocal images of secondary dendrites from DIV20 cultured rat hippocampal neurons transfected at DIV12 with either Scrambled or Sh-TSPAN5 constructs, both co-expressing GFP. After binding with α-GluA2 antibody, neurons were incubated for 0, 5 or 10 min. Secondary antibody was applied in non-permeabilising conditions for the extracellularly retained antibody (Extra, magenta) and in permeabilising condition for the internalised one (Intra, cyan). Scale bar = 5 μm. Right panel: quantification of GluA2 intracellular level normalised on total level (intracellular + extracellular). (GluA2 intracellular/total mean intensity: Scrambled 0 min 0.18±0.01, Scrambled 5 min 0.22±0.01, Scrambled 10 min 0.20±0.01, Sh-TSPAN5 0 min 0.16±0.01, Sh-TSPAN5 5 min 0.20±0.02, Sh-TSPAN5 10 min 0.21±0.01). n = Scrambled 0 min, 14; Scrambled 5 min, 14; Scrambled 10 min, 16 Sh-TSPAN5 0 min, 16; Sh-TSPAN5 5 min, 14; Sh-TSPAN5 10 min, 14 neurons. Values represent the mean ± SEM. * = p<0.05, ** = p<0.01, *** = p<0.001.

Another tetraspanin, TSPAN7, was previously shown to act on AMPAR internalisation (Bassani et al., 2012). To explore whether TSPAN5 could have a similar role, we evaluated the internalisation of AMPARs using an antibody-feeding paradigm. Rat hippocampal neurons transfected at DIV12 with either Scrambled or Sh-TSPAN5, were exposed to an α-GluA2 antibody directed against a surface epitope and incubated for different time points (Fig 4D). Both Scrambled and Sh-TSPAN5 transfected neurons exhibited a significant increase in the GluA2 intracellular/total ratio after 5 min, suggesting that AMPAR internalisation is not affected by TSPAN5 knockdown. Interestingly, in Scrambled-transfected neurons, recycling of the receptor at 10 min post incubation brought GluA2 back to the surface, with levels of the intracellular/total ratio similar to those at time-point 0; by contrast, the Sh-TSPAN5-transfected neurons maintained higher levels of internalised receptor at the 10 min time point. This potentially points to defects in GluA2 exocytosis.

### TSPAN5 regulates exocytosis of newly synthesised GluA2 subunits

Given the possible localisation of the TSPAN5, AP-4, Stargazin and AMPAR complex in Rab11 positive organelles, which have been shown to mediate receptor recycling back to the plasma membrane, we decided to directly evaluate GluA2 recycling. To this end, we relied on an overexpression model as recycling levels of endogenous GluA2 receptors are too low to be detected by an antibody feeding approach. We decided to use super-ecliptic pHluorin (SEP)-tagged GluA2, where the SEP has been inserted in the extracellular domain of GluA2 (Ashby et al., 2004; Hildick et al., 2012; Sankaranarayanan et al., 2000). SEP is extremely useful for intracellular trafficking studies as it is only fluorescent at a neutral pH, allowing for the visualisation of the receptor only when it is exposed to the neutral extracellular environment. Instead, its fluorescence is quenched in the acidic intracellular vesicles. We verified the functionality of the SEP-GluA2 by exposure to imaging media at pH 6, which completely abolished the signal, or to media containing NH_4_Cl that alkalinise also intracellular vesicles (Supplementary Figure related to Figure 5). To avoid interferences from endoplasmic reticulum (ER)-contained GluA2, we only evaluated the signal in individual dendritic spines, which are virtually devoid of ER (Rathje et al., 2013, 2014; Wilkinson et al., 2014). We applied a FRAP-FLIP protocol in which a portion of dendrite (ROI) is bleached and then imaged over 300 sec while repetitively bleaching the flanking regions of the ROI to eliminate interference from receptors laterally diffusing into the ROI (Hildick et al., 2012) (Fig 5A). This protocol allows for the selective visualisation of receptors that were present in intracellular compartments at the time of the initial bleaching. These receptors are quenched and would therefore not be affected by bleaching, retaining the ability to fluoresce once exposed to the neutral extracellular environment. As such, the protocol allows for the visualisation of newly synthesised receptors and internalised receptors recycling back to the plasma membrane. Application of cycloheximide before the experiment allowed us to block synthesis of new receptors, thereby restricting the analysis to recycling receptors only (Fig 5A). To our surprise, there was no difference in the levels or kinetics of SEP-GluA2 recycling upon modulation of TSPAN5 expression levels, obtained by the concomitant transfection of Scrambled, Sh-TSPAN5 or Rescue mCherry constructs (Fig 5B-D). Although the presence of cycloheximide could be blocking the synthesis of other proteins necessary for TSPAN5-dependent recycling of AMPARs, thus masking the effect of TSPAN5 knockdown, the most likely explanation for the reduction in surface GluA2 is that there could be a defect in the exocytosis of newly synthesised receptors. We thus used the same FRAP-FLIP approach but without application of cycloheximide; this experimental setup allows for the simultaneous observation of recycling receptor and exocytosis of newly synthesised receptor (Fig 5E). In this experiment, we observed a significant reduction of the recovery after photobleaching in individual dendritic spines of Sh-TSPAN5-transfected neurons compared to Scrambled-transfected neurons (Fig 5F-H). This defect was completely reversed in Rescue-transfected neurons, even showing a potentiation of the recovery. Although we cannot exclude a limited contribution of defects in GluA2 recycling, this was previously shown to participate less than 20-30% of the total exocytosis (Passafaro et al., 2001). Instead, upon TSPAN5 knockdown, the defect observed in absence of cycloheximide is roughly a 50% reduction, demonstrating that exocytosis of newly synthesised GluA2 receptors is regulated by TSPAN5.

**Figure 5.**
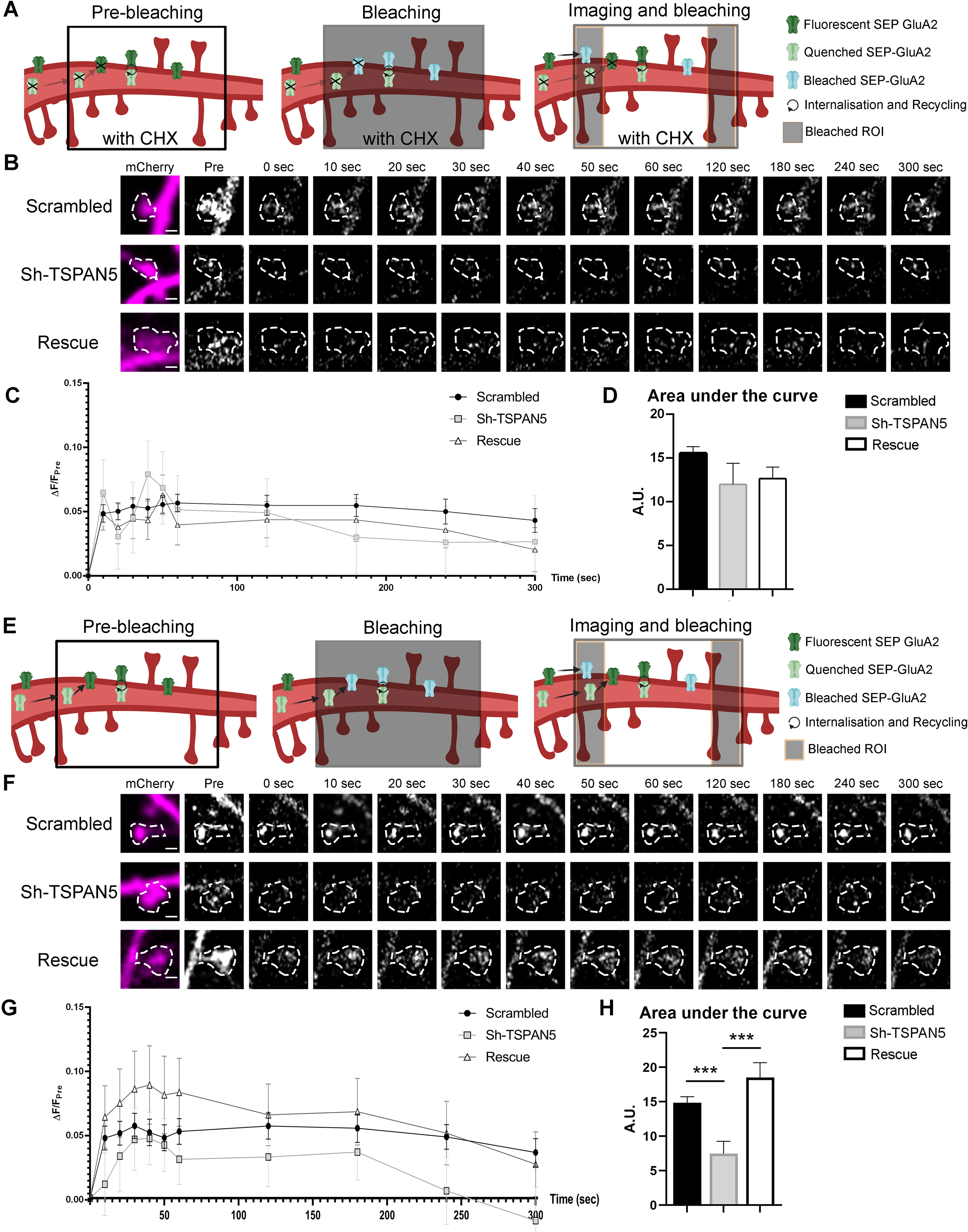
TSPAN5 regulates exocytosis of newly synthesised GluA2. **(A)** Schematic of the FRAP-FLIP experiment presented in panel B. SEP-GluA2 in pre-bleaching condition are either fluorescent (green) if exposed to the extracellular media or quenched (light green) if in intracellular compartments. A region of the dendrite is bleached (black box). SEP-GluA2 that were fluorescent (and so extracellularly exposed) at the time of bleaching becomes bleached (light blue). Quenched SEP-GluA2 are not affected by the bleaching. During imaging the ROI flanking regions are continuously bleached (black lateral boxes), thus lateral diffusing SEP-GluA2 will be bleached. Receptors that had been internalised and directed for recycling would be exocytosed and become fluorescent. Newly synthesised receptors would not be present due to the application of cycloheximide (CHX) (crossed out receptors). Controls for pH sensitivity of the SEP signal are shown in the supplementary figure relative to Figure 5. **(B)** Live confocal images of individual dendritic spines from DIV20 cultured rat hippocampal neurons transfected at DIV12 with SEP-GluA2 and either Scrambled, Sh-TSPAN5 or Rescue construct co-expressing mCherry. Neurons were treated for 2 h with 200 μg/ml of cycloheximide to inhibit protein synthesis and then imaged under a FRAP-FLIP protocol for 5 min to isolate the recycling receptors. mCherry (magenta) and SEP-GluA2 (white) images (timepoints: Prebleach, postbleach, 10, 20, 30, 60, 120, 180, 240 and 300 s) are shown. The dendritic spine mask is depicted with white dashed line. Scale bar = 1μm. **(C)** Quantification of the ΔF/F_pre_ for SEP-GluA2 over time for Scrambled-, Sh-TSPAN5- and Rescue-transfected neurons. **(D)** Quantification of the area under the curve relative to panel B (Area under the curve (A.U.): Scrambled 15.56±0.74, Sh-TSPAN5 11.99±2.51, Rescue 11.77±1.31). n = Scrambled, 56; Sh-TSPAN5, 53; Rescue, 53 dendritic spines. **(E)** Schematic of the FRAP-FLIP experiment presented in panel F. SEP-GluA2 at basal condition are either fluorescent (green) if exposed to the extracellular media or quenched (light green) if in intracellular compartments. A region of the dendrite is bleached (black box). SEP-GluA2 that were fluorescent (and so extracellularly exposed) at the time of bleaching becomes bleached (light blue). Quenched SEP-GluA2 is not affected by the bleaching. During imaging the ROI flanking regions are continuously bleached (black box), thus lateral diffusing SEP-GluA2 will be bleached. Receptors that have been internalised and directed for recycling would exocytose and become fluorescent. Newly synthesised receptors could also travel in intracellular vesicles to be exocytosed and become fluorescent. **(F)** Confocal images of individual dendritic spines from DIV20 cultured rat hippocampal neurons transfected at DIV12 with SEP-GluA2 and either Scrambled, Sh-TSPAN5 or Rescue construct co-expressing mCherry. Neurons were imaged under a FRAP-FLIP protocol for 5 min to analyse receptor exocytosis. mCherry (magenta) and SEP-GluA2 (white) images (timepoints: Prebleach, postbleach, 10, 20, 30, 60, 120, 180, 240 and 300 s) are shown. The dendritic spine mask is depicted with white dashed line. Scale bar = 1 μm. **(G)** Quantification of the ΔF/F_pre_ for SEP-GluA2 over time for Scrambled-, Sh-TSPAN5- and Rescue-transfected neurons. **(H)** Quantification of the area under the curve relative to panel E (Area under the curve (A.U.): Scrambled 14.85±0.89, Sh-TSPAN5 7.49±1.77, Rescue 18.5±2.18). n = Scrambled, 56; Sh-TSPAN5, 35; Rescue, 29 dendritic spines. Values represent the mean ± SEM. * = p<0.05, ** = p<0.01, *** = p<0.001.

### TSPAN5 regulates exocytosis of newly synthesised GluA2-containing AMPARs possibly by avoiding their degradation via the lysosomal pathway

To further confirm the role of TSPAN5 in AMPAR exocytosis, we took advantage of an endoplasmic reticulum (ER) retention system called ARIAD (Hangen et al., 2018; Rivera et al., 2000). In this system, the ARIAD-GluA2 is synthesised in the ER similarly to endogenous GluAs, but the presence of a conditional aggregation domain (CAD) results in its retention in this compartment. The protein can be released in a controlled manner by application of the ariad ligand that causes the disassembly of the CAD domains allowing the protein to continue along the secretory pathway. The fusion protein also presents a myc tag on the extracellular side allowing detection of the exocytosed receptor. As such, by applying an anti-myc antibody in the culture media after exposing the cells to the ariad ligand, one can assess the levels of plasma membrane inserted ARIAD-GluA2 directly coming from the ER site of synthesis (Fig 6A, B). As expected from our previous results, TSPAN5 knockdown resulted in a reduction in the surface levels of ARIAD-GluA2 90 min after application of the ariad ligand, an effect that was rescued by re-expression of the Sh-resistant form of TSPAN5 (Fig 6C). We also analysed dendritic transport of ARIAD-tdTomato-GluA2 via live imaging of neurons 30 min after addition of the ariad ligand. Here, we did not detect any change in the average speed of transport of GluA2 containing vesicles in either the anterograde or retrograde direction (Fig 6D, E), nor in the average number of vesicles (Fig 6F). These results suggest that there is either a lower amount of GluA2 loaded into vesicles directed for exocytosis or that these vesicles fail to deliver their content to the plasma membrane of dendrites and might be directed for degradation as a result.

**Figure 6:**
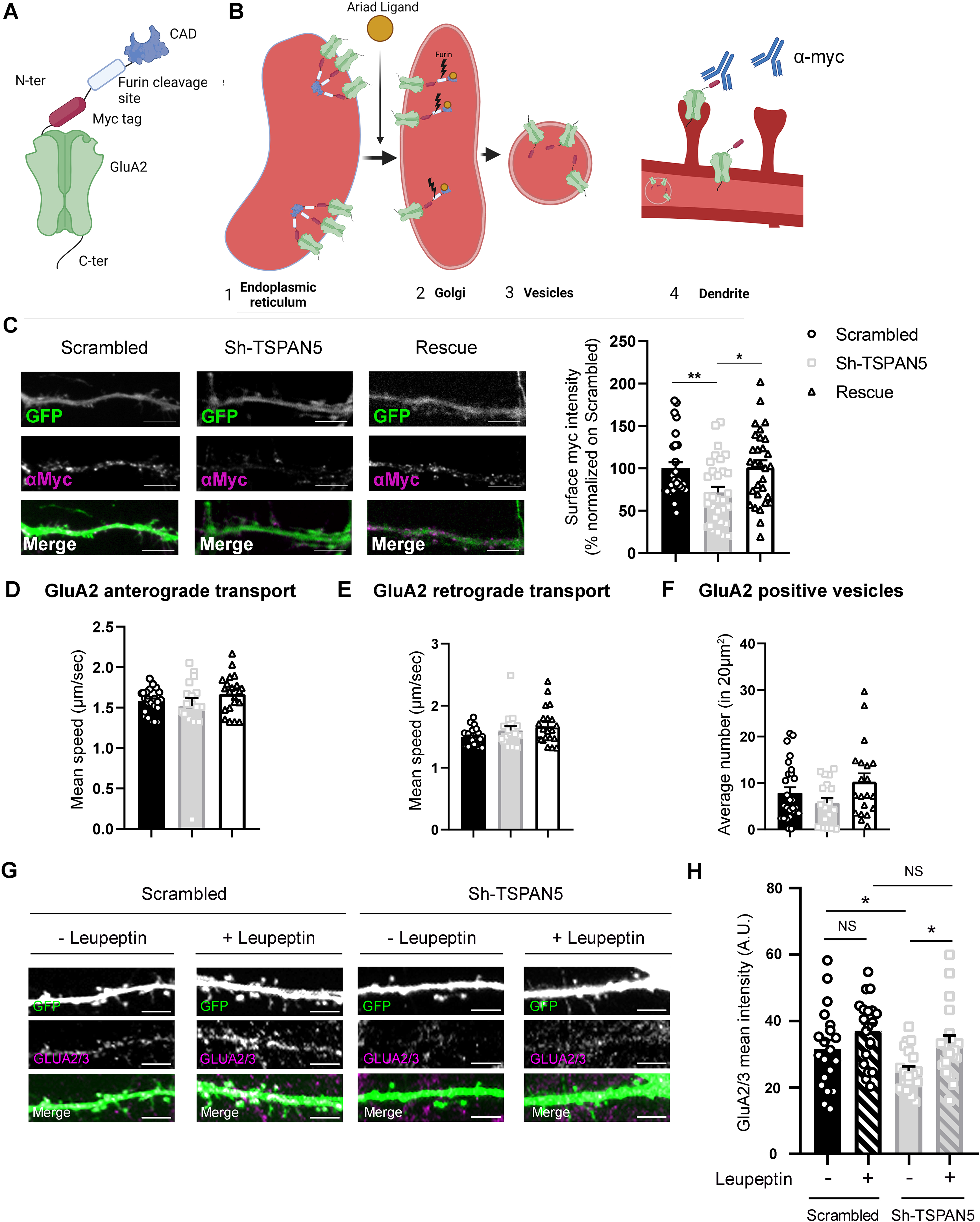
TSPAN5 regulates exocytosis of newly synthesised GluA2 containing AMPARs possibly by preventing their degradation via the lysosomal pathway. **(A)** Schematic of the ARIAD-GluA2 construct. **(B)** In basal conditions (1), ARIAD-GluA2 is retained in the endoplasmic reticulum (ER) due to the self-assembly properties of the CAD domain. Upon its application (2), the Ariad ligand binds to CAD, inhibits self-assembly and allows the ARIAD-GluA2 to move to the Golgi where the endogenous Furin protease cleaves the CAD domain. As such, ARIAD-GluA2 is free to be loaded onto secretory vesicles (3), transported along the dendrites and subsequently exocytosed (4). Application of an anti-myc antibody in the culture medium allows for the detection of the plasma membrane pool of GluA2 that was released from the ER after application of the Ariad ligand. **(C)** Left panel: Confocal images of DIV20 rat cultured hippocampal neurons transfected at DIV12 with the ARIAD-myc-GluA2 construct and with either plasmids coding for GFP (green) and Scrambled, Sh-TSPAN5 or Rescue, respectively, and immunostained with an anti-myc antibody in live staining conditions (magenta) 90 min after the application of the ariad ligand. Right panel: quantification of the surface anti-myc mean intensity normalised to Scrambled (Scrambled 100±7.14; Sh-TSPAN5 71.44±6.81; Rescue 101.4±7.92). n = 27-31 neurons per condition. Scale bar = 5 μm. **(D)** Quantification of the average speed of tdTomato positive organelles moving from the soma outwards along dendrites after application of the ariad ligand in DIV20 rat cultured hippocampal neurons transfected with ARIAD-tdTomato-GluA2 and with either Scrambled, Sh-TSPAN5 or Rescue. Mean speed (μm/sec) (Scrambled 1.58±0.03; Sh-TSPAN5 1.52±0.1; Rescue 1.67±0.05). **(E)** Quantification of the average speed of tdTomato positive organelles moving from the dendrites towards the soma after application of the ariad ligand in DIV20 rat cultured hippocampal neurons transfected with ARIAD-tdTomato-GluA2 and with either Scrambled, Sh-TSPAN5 or Rescue. Mean speed (μm/sec) (Scrambled 1.67±0.21; Sh-TSPAN5 1.81±0.22; Rescue 1.67±0.07). **(F)** Quantification of the average number of tdTomato positive vesicles in a 20 μm^2^ stretch of a dendrite after application of the ariad ligand in DIV20 rat cultured hippocampal neurons transfected with ARIAD-tdTomato-GluA2 and with either Scrambled, Sh-TSPAN5 or Rescue. Average number per 20 μm^2^ (Scrambled 7.9±1.2; Sh-TSPAN5 5.8±1.1; Rescue 10.3±1.8). **(G)** Confocal images of secondary dendrites from DIV20 rat cultured hippocampal neurons transfected at DIV12 with either Scrambled or Sh-TSPAN5 constructs, both co-expressing GFP. Neurons were treated for 90 min with either vehicle (H_2_O) or leupeptin (100 μM), fixed and immunostained for GLUA2/3 (magenta). Scale bar = 5 μm. **(H)** Relative quantification of GluA2/3 staining mean intensity (GluA2/3 mean intensity: Scrambled vehicle 31.25±2.43; Scrambled leupeptin: 36.95±2.25: Sh-TSPAN5 vehicle: 24.51±1.35; Sh-TSPAN5 leupeptin 33.3±2.22). n = Scrambled vehicle, 20; Scrambled leupeptin, 20; Sh-TSPAN5 vehicle, 20; Sh-TSPAN5 leupeptin, 20 neurons. Values represent the mean ± SEM. * = p<0.05, ** = p<0.01, *** = p<0.001.

To test this second possibility, we assessed the total levels of GluA2/3 via immunofluorescence in DIV20 neurons transfected at DIV12 with either Scrambled or Sh-TSPAN5 and treated with the lysosomal inhibitor leupeptin (Fig 6G), considering that AMPARs are mostly degraded via this pathway (Ehlers, 2000). Leupeptin treatment increased GluA2/3 to similar levels in Scrambled and Sh-TSPAN5 transfected neurons, supporting the hypothesis that AMPARs are degraded in the absence of TSPAN5 (Fig 6H). Altogether, our data support a model whereby the association of TSPAN5 with GluA2, occurring via AP-4 and Stargazin, promotes the exocytosis of newly synthesised GluA2-containing AMPARs, potentially via Rab11/TfR positive recycling endosomes (Fig 7).

**Figure 7.**
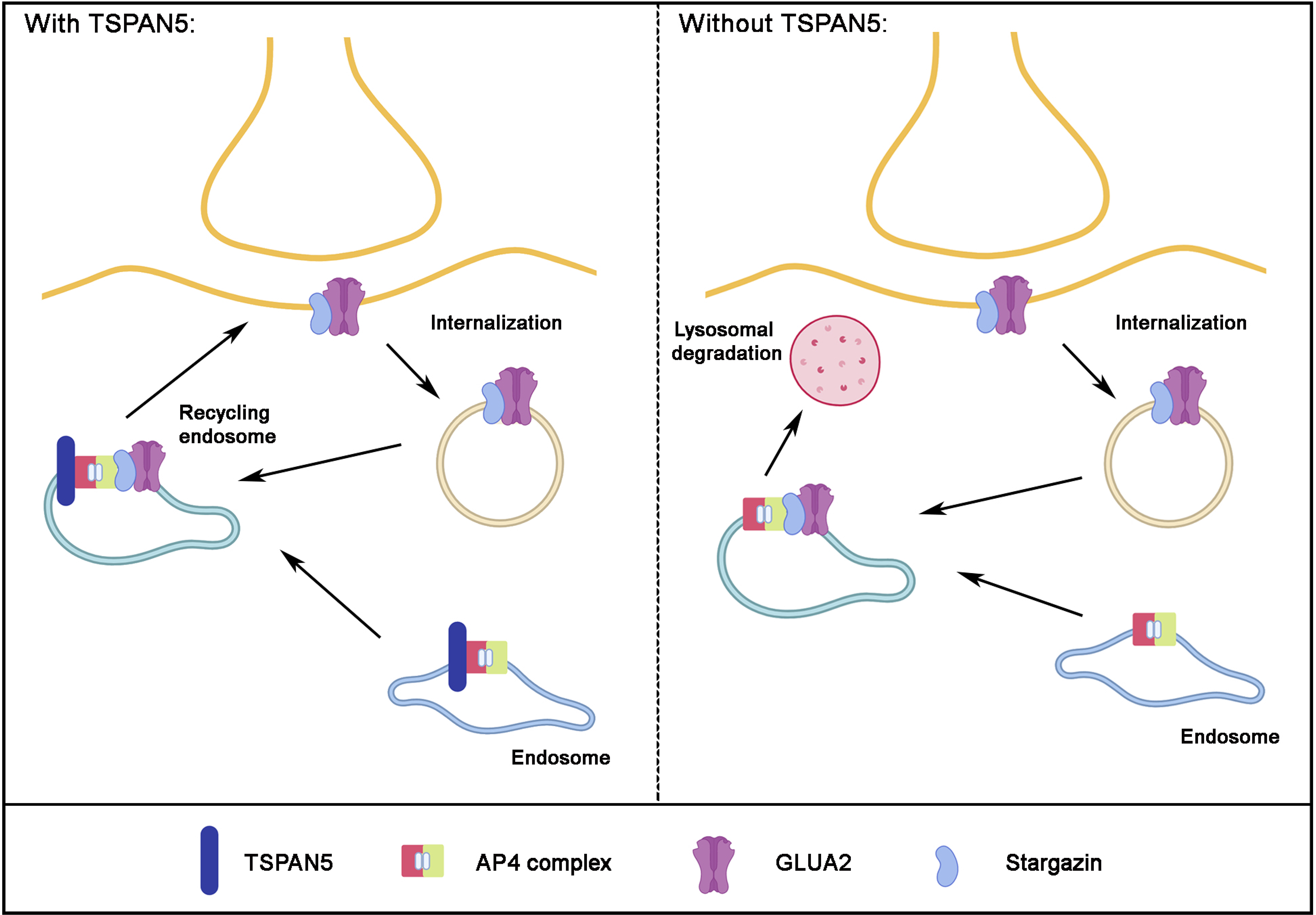
TSPAN5 regulates GluA2 exocytosis through recycling endosomes by the formation of a tetrameric complex with AP-4 and Stargazin. Schematic representation of TSPAN5 function in mature neurons (left) and TSPAN5 silencing effects (right). In our working model, TSPAN5 forms a complex with Stargazin and GluA2-containing AMPARs possibly in the endoplasmic reticulum or in a subsequent ERGIC vesicle. The presence of TSPAN5 is necessary to direct the GluA2 and Stargazin complex to the plasma membrane. TSPAN5 silencing in neurons induces the redirection of GluA2-containing vesicles to lysosomal degradation.

## Discussion

In this work, we have identified an intracellular pool of TSPAN5 that is enriched in mature hippocampal neurons. This pool participates in the delivery of newly synthesised GluA2-containing AMPARs to the plasma membrane. We showed that TSPAN5 forms a complex with AP-4, Stargazin and GluA2 and that this interaction could take place in recycling endosomes. Although the main function of recycling endosomes is to redirect endocytosed receptors back to the plasma membrane, they have also been shown to participate in a non-canonical secretory pathway. Proteins synthesised in the dendritic ER are trafficked to an ER-Golgi intermediate compartment (ERGIC) before being loaded to recycling endosomes for insertion in the plasma membrane (Bowen et al., 2017; Hirling, 2009). As a result, the receptors would bypass the Golgi compartment, which is poorly present in dendrites and dendritic spines. The molecular regulators of this process are not well defined. However, our data do not fully identify the nature of the organelles involved, therefore further investigation is required.

The differential effect of TSPAN5 knockdown on surface levels of GluA2 and GluA1 seems to indicate that GluA2- and GluA1-containing AMPARs follow different paths, although they can both interact with Stargazin (Chen et al., 2000). The trafficking of GluA2 and GluA1 has previously shown to be regulated by separate mechanisms for example, GluA2 delivery and recycling is a constitutive process, whereas GluA1 exocytosis to the plasma membrane is more dependent on synaptic plasticity (Passafaro et al., 2001; Shi et al., 2001). The observation that both surface and total levels of GluA1 are increased upon TSPAN5 knockdown suggests that compensatory mechanisms are exploited to maintain normal synaptic activity. This could also be a potentiation effect of a secretory pathway which does not rely on TSPAN5 and that is responsible for GluA1-containing AMPAR delivery. Our findings do not elucidate whether AMPARs exhibit a different subunit composition upon TSPAN5 knockdown. However, the fact that we observe a reduction in GluA2 and GluA3 and an increase in GluA1 potentially suggests that there could be an overall reduction in GluA2/3 tetramers and that the remaining GluA2 could potentially be redirected to GluA1/2 tetramers. In addition, an increase of GluA1 homomers could also occur. This could partially explain our previous observation that AMPAR-mediated mEPSCs are not affected by TSPAN5 knockdown in either their amplitude or frequency, but display altered kinetics (Moretto et al., 2019b) which can be due to a variation in the receptor subunit composition (Wei et al., 2009).

Our results also shed new light on AP-4 function. AP complexes select transmembrane proteins via interaction through typical sorting motifs and promote their insertion into specific vesicles (Robinson, 2004). A role for AP-4 in AMPARs intracellular trafficking was previously shown (Matsuda et al., 2008, 2013b): AP-4 was found to restrict AMPARs from being directed towards the axonal compartment. AP-4β knockout mice presented with mislocalisation of AMPARs to the axon, which accumulated in autophagosomes. The authors did not detect a reduction of dendritic AMPARs in AP-4 β knockout neurons, however this could have been due to compensatory mechanisms arising *in vivo* upon constitutive knockout of the AP-4 complex or because only levels of overexpressed AMPARs were analysed. It remains possible that other AP complexes could compensate for the loss of AP-4. In particular, AP-1 was found to regulate sorting and exocytosis of membrane proteins (Bonifacino, 2014b). Interestingly, the involvement of AP-4 in AMPAR exocytosis could potentially explain the intellectual disability phenotype of AP-4 deficiency syndrome; an imbalance between GluA2 and GluA1 subunits in the composition of AMPARs was previously shown to cause changes in how neurons respond to synaptic plasticity events, thus impacting on learning and memory functions (Moretto et al., 2017).

Together with our previous work, these data highlight the importance of TSPAN5 for neuronal function. TSPAN5 appears to act on two independent pathways; on the one hand, its localisation at the plasma membrane is crucial for the maturation of dendritic spines during neuronal development (Moretto et al., 2019a); on the other hand, TSPAN5 localisation in intracellular vesicles in mature neurons regulates exocytosis of GluA2-containing AMPARs enabling correct synaptic functionality.

## Acknowledgments

We sincerely thank Skye Stuart for critical reading of the manuscript. We thank Robert Malinow for the pCI-SEP GluR2 (Addgene plasmid # 24001). We thank Michisuke Yuzaki for the GST-Ct-Stargazin plasmid. We thank Margaret Robinson for the anti AP-4σ antibody. We thank Cecilia Gotti for the anti GluA2/3 antibody.

The financial support of Fondazione Telethon, Italy (GGP17283) is gratefully acknowledged. Part of this work was supported by PRIN (Progetti di rilevante interesse nazionale – Bando 2017), 20172C9HLW and Fondazione Cariplo, Italy (2019-3438).

## Funding

Fondazione Telethon, Italy (GGP17283)

PRIN (Progetti di ricerca di rilevante interesse nazionale – Bando 2017), 20172C9HLW

Fondazione Cariplo, Italy (2019-3438)

## Declaration of interests

The authors declare no competing interests.

## Author contribution

E.M. and M.P. conceptualised the project; E.M., A.L., F.M., C.B., F.C., F.J. and L.C. performed the investigation, formal analysis and wrote the manuscript. M.P. supervised, acquired funding and reviewed and edited the manuscript.

## EXPERIMENTAL MODELS

Animal procedures were performed in accordance with the European Community Council Directive of November 24, 1986 (86/609/EEC) on the care and use of animals. Animal procedures were approved by the Italian Ministry of Health (Protocol Number N° 2D46AN.463).

The HEK293FT cell line used to generate the lentiviruses was grown in DMEM supplemented with 10% FBS, 1% L-glutamine, 0.1% gentamycin and 0.1% G418 (Invitrogen). N2A cells were cultured in DMEM supplemented with 10% FBS, 2mM L-glutamine, 100 U/ml penicillin and 0.1 mg/ml streptomycin (Invitrogen).

Primary hippocampal neurons were prepared from Wistar E18 rat brains (Folci et al., 2014; Valnegri et al., 2011; Zapata et al., 2017). Neurons were plated onto coverslips coated overnight with 0.25 mg ml^−1^ poly-D-lysine (Sigma Aldrich) at 75,000 per well and grown in Neurobasal medium supplemented with 2% B27 (prepared as in (Chen et al., 2008), 0.25% L-glutamine, 1% penicillin/streptomycin and 0.125% Glutamate (Sigma Aldrich).

3-month-old male Wistar rats were used for hippocampus and cortex lysates.

## METHOD DETAILS

### Plasmids

pLVTHM-Scrambled, pLVTHM-Sh-TSPAN5, pSicor-TSPAN5-GFP (Rescue), pSicor-Scrambled-mCherry, pSicor-Sh-TSPAN5-mCherry, pSicor-TSPAN5-mCherry and pGEX4T1-TSPAN5-Ct have been characterised in our previous work (Moretto et al., 2019b). Rab4-GFP, Rab7-GFP and Rab11-GFP are kind gifts from Prof G. Schiavo. pCl-SEP-GluA2 was obtained from Addgene #24001 (Kopec et al., 2006). The pLenti-U6-(BsmBI)-hSyn-SaCas9-P2A-EGFP vector allowing the expression of *Staphylococcus aureus* Cas9 and a guide RNA for the knockdown of AP4 β and AP4 ε were constructed by replacing the EF-1α promoter in the pLenti_SaCRISPR-EGFP plasmid (gift from Christopher Vakoc; Addgene 118636) with the hSyn promoter from the pAAV-hSyn-EGFP plasmid (gift from Bryan Roth; Addgene 50465). The gRNA sequences were inserted downstream of the U6 promoter using BsmbI cloning sites. EGFP expression was used for visualisation of the transduced neurons. The guide RNAs were CCGGTAGCGCAGCCTATCAGC and TTGATGAATCCTTACGAAGAG for AP4 β and ε, respectively. The control non-targeting guide RNA sequence was GTTCCGCGTTACATAACTTA.

### Yeast Two-Hybrid Screening

For Yeast two-hybrid experiments, a fragment corresponding to the TSPAN5 C-terminal tail (aa 254-268) was cloned in frame with the GAL4 DNA-binding domain (pGBKT7 vector) and used as bait to screen a human adult brain cDNA library (Clonetech, Mate and Plate Library). Positives clones (3+) grew on plates containing X-α-GAL and Aureobasidin A (QDO/X/A plates) and expressed all four integrated reporter genes: HIS3, ADE2, AUR1C and MEL1 under the control of three distinct Gal4-responsive promoters. cDNA plasmids from positive clones were recovered via DH5a *Escherichia coli* (*E. coli*) and sequenced.

### Transfection and Infection

For lentivirus production, HEK293FT cells were transfected using the calcium phosphate method. Briefly, DNA was mixed with 130mM CaCl_2_ in H_2_0. One volume of HEBS buffer (280 mM NaCl, 100 mM Hepes, 1.5 mM Na2HPO4, pH 7.11) was added to the DNA and thoroughly mixed to produce air bubbles. The mix was added to the cells and left for 5 h before washing and changing the medium.

Transfection of N2a cell was performed in 60-70% confluent cultures seeded in 6-well plates at 200,000 cells/well the previous day. Cells were transfected with 3 μg DNA/well using the calcium phosphate method and used 48 hours post-transfection.

Rat hippocampal neurons were transfected with Lipofectamine 2000 (Invitrogen) following the manufacturer’s instructions or infected with lentiviral particles produced as previously described (Lois et al., 2002).

### BS3 crosslinking

Experiments were carried out according to (Boudreau et al., 2012). Briefly, primary hippocampal neurons were washed twice with PBS supplemented with 0.1 mM CaCl_2_ (Sigma Aldrich) and 1 mM MgCl_2_ (Sigma Aldrich) at 37°C. Neurons were then exposed to PBS supplemented with 0.1 mM CaCl_2_ and 1 mM MgCl_2_ with or without the BS3 crosslinker (1 mg/ml, ThermoFisher) at 4°C for 10 min. Neurons were then rapidly washed first with TBS supplemented with 0.1 mM CaCl_2_ and 1 mM MgCl_2_ plus 50 mM glycine (Sigma Aldrich) at 4°C and subsequently with TBS supplemented with 0.1 mM CaCl_2_ and 1 mM MgCl_2_ at 4°C prior to lysis with BS3 buffer (50 mM Tris-HCl, 150 mM NaCl, 1 mM EDTA, pH 7.4, 1% SDS plus protease inhibitors). 3X Laemmli sample buffer was then added and samples were analysed by SDS-PAGE and western blotting. Crosslinked proteins present in the plasma membrane appeared as high molecular bands. All the other bands, which were also present in the non-crosslinked reaction were considered as part of the intracellular pool. Extracellular and intracellular intensities were normalised on tubulin intensity.

### Vesicles purification

Hippocampi and cortices were collected from adult Wistar rats and homogenised with glass-teflon potter in homogenisation buffer (0.32 M sucrose, 20 mM Hepes-NaOH, protease inhibitors, pH 7.4). The total homogenate was centrifuged at 1,000 g for 10 min at 4°C. The supernatant S1 was further centrifuged at 10,000 g for 15 min at 4°C. The resulting pellet, corresponding to crude synaptosomal fraction, was resuspended in homogenisation buffer and centrifuged again at 10,000 g for 15 min at 4°C to wash the synaptosomes. Crude synaptosomes were lysed using hypotonic shock. The resulting vesicles were layered on a 9 ml 50–1000 mM sucrose gradient and centrifuged in a SW40Ti Beckman rotor at 65,000 g for 3 h. After centrifugation, 10 equal fractions were collected from the top of the gradient, and protein precipitation was performed using 6% trichloroacetic acid (TCA) and 0.02% deoxycholate. 3X sample buffer was then added and the samples analysed by SDS-PAGE and western blot.

### Immunoprecipitation

For immunoprecipitation experiments, hippocampi and cortices were dissected from adult rat brains, pooled together and lysed in RIPA buffer (50 mM Tris, 150 mM NaCl, 1 mM EDTA, 1% NP40, 1% Triton X-100, pH 7.4, protease inhibitor) with a tephlon-glass potter, rotated for 1 h at 4°C and centrifuged at 10,000 g for 30 min at 4 °C. Supernatants were incubated with antibodies at 4 °C overnight. Protein A-agarose beads (GE Healthcare, USA) were incubated with the supernatant at 4 °C for 2 h. Beads were washed three times with RIPA buffer, resuspended in 3X Laemmli sample buffer and analysed by SDS–PAGE followed by western blotting.

### GST pulldown

GST-fusion proteins were prepared by growing transformed BL21 *E. coli* and inducing recombinant protein expression by adding IPTG (0.5mM final concentration) for 2 h. Bacteria was pelleted, and the GST-fusion protein was purified employing standard procedures using glutathione Sepharose beads (Thermo Scientific).

Hippocampi and cortices dissected from adult rat brains were pooled together, lysed in RIPA buffer by homogenisation in a tephlon-glass potter, rotated for 1 h at 4°C and then centrifuged at 10,000 g for 30 min at 4 °C. Supernatants were incubated with glutathione sepharose beads for 3 h at 4°C and then washed and resuspended in 3X sample buffer and analysed by SDS-PAGE followed by western blotting.

### Western blots

Proteins were transferred from the acrylamide gel onto the nitrocellulose membrane (0.22 μm, GE Healthcare). Membranes were incubated with the primary antibodies (α-TSPAN5, Aviva System biology #AV46640, 1:500; α-transferrin receptor, Thermo Fisher Clone H68.4, 1:500; α-tubulin, Sigma Aldrich T5168, 1:40,000; α-AP-4σ, gift from Dr Margaret Robinson, 1:500; α-AP4-ε, BD biosciences 612018, 1:1000; α-GluA1, Cell Signalling #13185, 1:1000; α-GluA2/3, gift from Dr Cecilia Gotti, 1:2,000; α-Stargazin, Cell Signalling #8511, 1:1000; α-EEA1, BD transduction laboratories Clone 14, 1:2000; α-Rab11 BD transduction laboratories Clone 47, 1:1500; α-Rab7, SySy 320003, 1:700; α-VGlut1, SySy 135303, 1:2000; α-GFP, MBL #598, 1:2,500; α-NR2A, Neuromab N327/95, 1:1000; α-CD81, Santa Cruz Biotech #166029, 1:1000) at room temperature for 2–3 h or overnight at 4°C in TBS Tween-20 (0.1%), milk (5%). After washing, the blots were incubated at room temperature for 1 h with horseradish peroxidase-conjugated α-rabbit, α-mouse or α-rat antibodies (1: 2,000) in TBS Tween-20 (0.1%), milk (5%). Immunoreactive bands on blots were visualised by enhanced chemiluminescence (Chemidoc XRS+, BioRad).

### Immunocytochemistry

Cultured hippocampal neurons were washed in PBS supplemented with 0.1 mM CaCl_2_ and 1 mM MgCl_2_ and fixed in paraformaldehyde (4%, Sigma Aldrich)/sucrose (4%, Sigma Aldrich) for 10 min at room temperature and incubated with primary antibodies (α-TSPAN5, Aviva System Biology #AV46640, 1:50; α-GluA2/3, gift of Dr Cecilia Gotti, 1:500) in GDB1X solution (2X: 0.2% gelatin, 0.6% Triton X-100, 33mM Na_2_HPO_4_, 0.9 M NaCl, pH 7.4) for 2 h at room temperature.

For surface staining, antibodies (α-GluA2, Merck clone 6C4, 1:200; α-GluA1, Cell signaling #13185, 1:150, α-myc, Sigma #M5546, 1:1000) were applied to neurons for 10 min at room temperature followed by a washing step in PBS supplemented with 0.1 mM CaCl_2_ and 1 mM MgCl_2_ and paraformaldehyde fixation.

After three washes with high salt buffer (500mM NaCl, 20mM NaPO_4_^2-^, pH 7.4), coverslips were incubated with secondary antibodies (Alexa-conjugated: 1:400; DyLight-conjugated: 1:300) in GDB1X solution for 1 h at room temperature.

Internalisation experiments were performed as described by Bassani and colleagues (Bassani et al., 2012). Briefly, neurons were incubated with the anti-GluA2 surface epitope antibody at 10 μg/ml in culture medium for 10 min at room temperature. Excess antibody was then removed by washing with PBS c/m. The antibody-bound receptors were then allowed to undergo internalisation for 0, 5 or 10 min in the original media at 37°C. After paraformaldehyde fixation, a secondary antibody labelled with AlexaFluor 555 was incubated in non-permeabilising condition (PBS supplemented with 10% goat serum) for 1 h at room temperature. After washing, the coverslips were incubated with a secondary antibody labelled with DyeLight-649 in permeabilising condition (GDB1X) for 1 h at room temperature.

Coverslips were washed with high salt buffer and mounted with Mowiol (Sigma Aldrich). Fluorescence images were acquired with an LSM800 Meta confocal microscope (Carl Zeiss) and a 63X oil-immersion objective (numerical aperture 1.4) with sequential acquisition settings, at 1,024 × 1,024 pixels resolution. Images were collected as Z-stack series projections of approximately 6–10 images, each averaged four times and taken at depth intervals of 0.75 μm.

Dendritic spines were counted on all GFP-positive neuronal dendritic arbor excluding the soma and classified with NeuronStudio software (NeuronStudio©) according to the following parameters: General parameters for spine identification: Length > 0.2 μm and < 3.0 μm, Max Width 3.0 μm, Stubby spines size > 10 voxels, Non Stubby spines size > 5 voxels. For spine type classifications, the following logical tests were used: if Neck Ratio (head/neck diameter) > 1.100 then a spine was classified as Thin (if also spine length/head diameter > 2.5) or Mushroom (if also head diameter was > 0.35 μm). A spine is classified as Stubby if it fails any of the precedent logical tests.

For quantification, a mask was drawn on the GFP or mCherry channel and the immunofluorescence signal for the different antibodies was quantified as mean intensity. For the analysis of surface GluA2 or GluA1 in Fig 3, dendritic spine regions were identified via NeuronStudio as stated above and the quantification performed only on the corresponding areas.

### FRAP-FLIP imaging of SEP-GluA2

Neurons transfected with pCl-SEP-GluA2 and either Scrambled, Sh-TSPAN5 or Rescue mCherry constructs were incubated for 15 min in equilibrated Tyrode’s buffer (15 mM D-glucose, 108 mM NaCl, 5 mM KCl, 2 mM MgCl2, 2 mM CaCl2 and 25 mM HEPES-NaOH, pH 7.4) and coverslips were mounted in an open Inox chamber (Life imaging services). For recycling only experiments, neurons were previously incubated with 200 μg/ml Cycloheximide (Life Technologies) for 2 hr with cycloheximide also present in the recording Tyrode’s buffer. A LSM800 confocal microscope equipped with an environmental chamber (37°C, 5% CO2) was used. A secondary dendrite from neurons positive for both mCherry and SEP signal was selected and a portion of the dendrite (ROI) was initially bleached with high 488nm laser power (80%) and then sequentially bleached at the extremities of the ROI and imaged every 500 ms. The fluorescence intensity of SEP-GluA2 on individual dendritic spines was measured for individual time points and normalised as F_n_-F_0_ (ΔF)/F_prebleach_. The area under the curve was measured via GraphPad Prism 8 as area below a curve fitted by regression on the average values.

### Real-time PCR

mRNA was extracted from N2A cells or cultured rat hippocampal neurons using Nucleozol Reagent following the manufacturer instructions (Macherey Nagel).

For each condition, 1.5 μg of extracted mRNA was used to synthetise cDNA using SuperScript VILO cDNA Synthesis Kit (Thermo Fisher).

The target sequences of AP4B, AP4E and β-actin (endogenous control) were amplified from 60 ng of cDNA in the presence of SYBR Green PCR Master Mix (Applied Biosystems) using Applied Biosystems 7000 Real-Time thermocycler. Primer sequences were as follows: AP4B Fw (AGTTGCTGGGACTTCGACAA), AP4B Rv (CCGTGGACCCCAAGTAACC), AP4E Fw (TTCTGGATGGTTTTGTGGCTG), AP4E Rv (CCAGTGAAGCCAGATGAAGAAAA), β -actin Fw (AGATGACCCAGATCATGTTTGAGA), β -actin Rev (CCTCGTAGATGGGCACAGTGT) Each sample was run in triplicate, and results were calculated using the ΔΔCT method to allow normalisation of each sample to the internal standard and comparison with the calibrator of each experiment.

### Experiments with ARIAD constructs

90 min after addition of the ariad ligand (2 μM), anti-myc antibody (Sigma, #M5546, 1:1000) was added to the media. Neurons were directly fixed with 4% PFA, 4% sucrose and then incubated with a secondary antibody anti-mouse Alexa 565. Images were taken with a Leica DM5000 microscope with a 40X objective. Quantification was performed with ImageJ to quantify the surface receptor mean intensity.

Intracellular transport videos were acquired on an inverted Leica microscope (DMI6000B) at the Bordeaux Imaging centre at DIV18-19. This microscope, controlled with Metamorph (Molecular devices, Sunnyvale, USA) is equipped with a confocal spinning-disk system (Yokogawa CSU-X1, laser: 491nm, 561nm), an EMCCD camera (Photometrics Quantem 512), a FRAP scanner (Roper Scientific, Evry, France, 561nm) and an oil objective HCX PL Apo 100X 1.4 NA. The coverslips were mounted in a Ludin chamber with 1 ml of Tyrode medium (15 mM glucose, 100 mM NaCl, 5 mM KCl, 2 mM MgCl_2_, 2 mM CaCl_2_, 10 mM HEPES, 247 mosm/l) with 2 μM of ARIAD ligand to release the proteins of interest from the ER, and placed at 37°C in a Life Imaging Services chamber. Videos were acquired between 30 and 60 min of incubation with the ligand using the following acquisition sequence (Hangen et al. 2018): 10 images are acquired (100ms exposure), followed by the photobleaching of ~60 μm^2^ of proximal dendrite (5 repetitions, 70% laser), followed by video acquisition (1 min at 1 Hz, 300 images, 100 ms exposure). Co-transfection with the sh-RNAs or control were confirmed by the acquisition of an image in the green channel (488nm) prior to the video recording.

The videos were analysed by generating kymographs thanks to the ImageJ plugin KymoToolBox (Hangen et al. 2018). The vesicles’ pathways were traced by the deep learning software KymoButler (Jakobs et al. 2019). From those traces, the number of vesicles and mean speed were calculated.

### Schematic figure

The schematics in Figure 5A, 5E, 6A, 6B and 7 were prepared using BioRender software (Biorender.com).

### Quantification

All statistical analyses were done with GraphPad Prism 8 software.

Two-tailed unpaired t-test was performed to assess statistical significance between two independent groups (Fig 1A). One-way ANOVA, followed by Newman-Kuls post-hoc multiple comparison test, was used to assess statistical significance between three or more groups (Fig 1B, 2D, 3A, 3B, 3D, 4A, 4B, 4C, 4D, 5D, 5G, 6A, 6B, 6C, 6D, 6F).

Statistical details of the experiments can be found in the figure legends (exact mean values, standard errors of the mean (SEM) and n).

Western blots were repeated at least three times from three independent experiments. Imaging experiments on cultured neurons were performed on at least three independent cultures.

## Supplementary figure legends

**Supplementary figure related to Figure 1:**
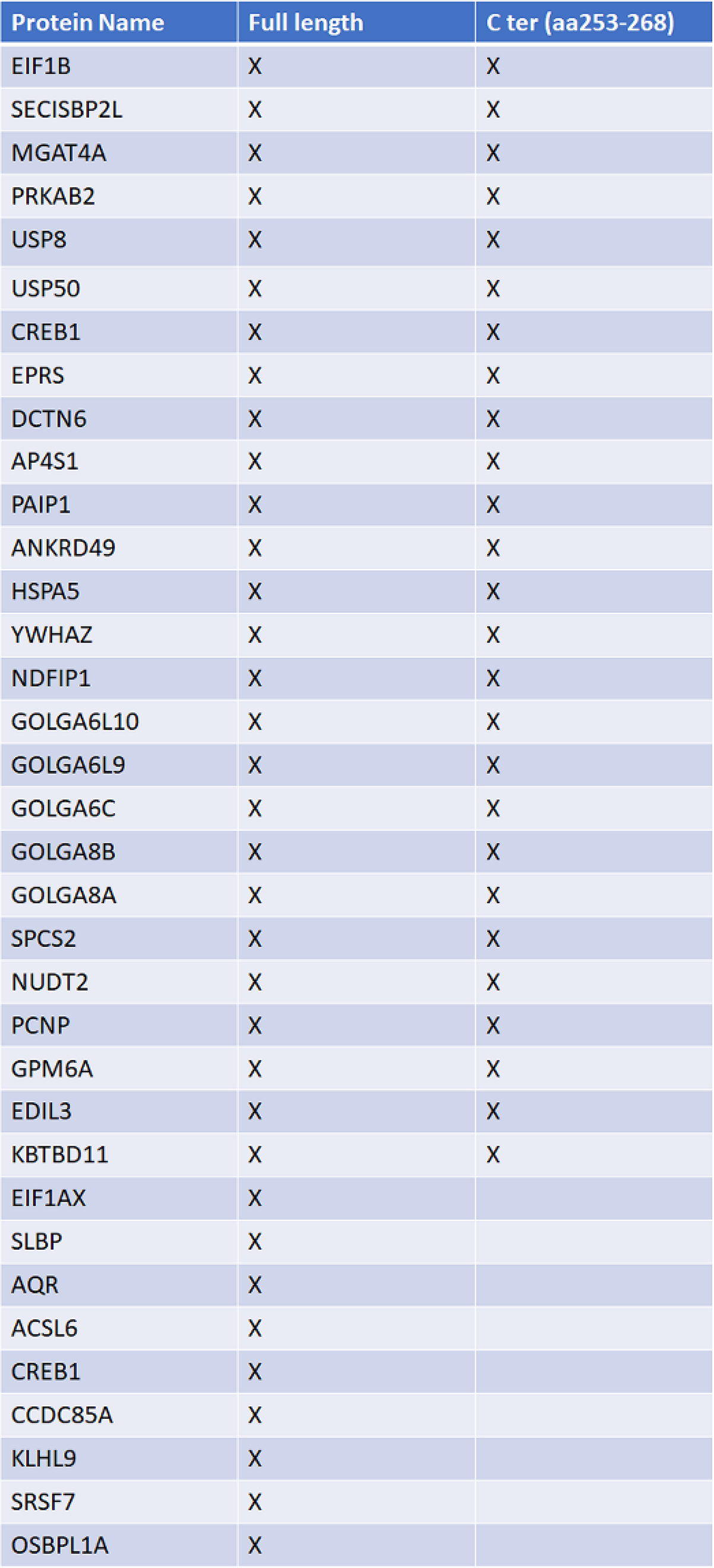
List of proteins identified in the two-hybrid screen performed using either the full-length TSPAN5 protein or its C-terminus only.

**Supplementary figure related to Figure 3:**
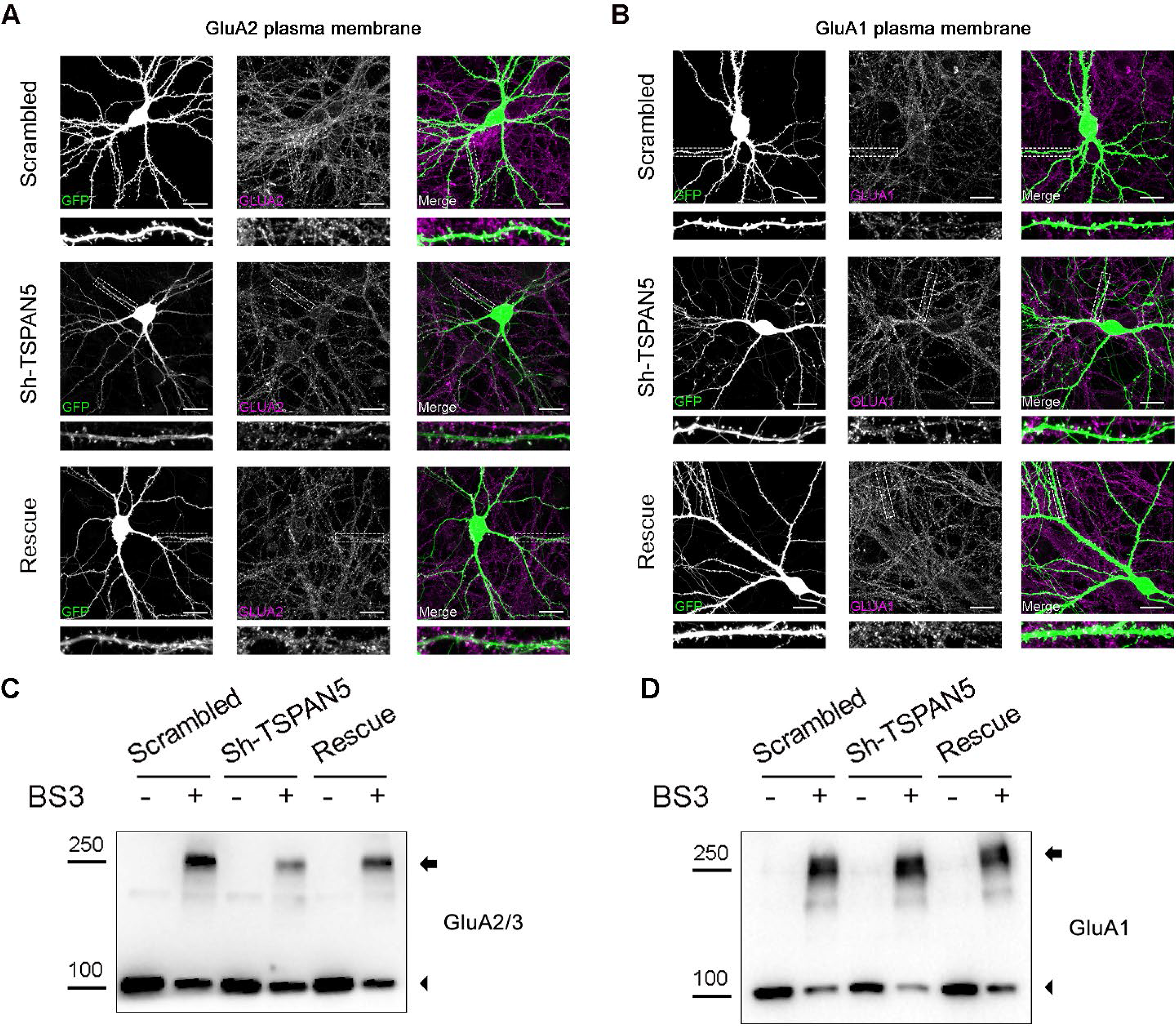
**(A)** Confocal images of dendrites from cultured rat hippocampal neurons transfected at DIV12 with either Scrambled, Sh-TSPAN5 or Rescue constructs, all co-expressing GFP (green) and immunostained at DIV20 with an antibody against an extracellular epitope of GluA2 in non-permeabilised conditions (magenta). Inserts show higher magnification of the dendrites highlighted with a white dashed line, which are presented in Figure 3A. Scale bar = 20 μm. **(B)** Confocal images of dendrites from cultured rat hippocampal neurons transfected at DIV12 with either Scrambled, Sh-TSPAN5 or Rescue constructs all co-expressing GFP (green) and immunostained at DIV20 with an antibody against an extracellular epitope of GluA1 in non-permeabilised conditions (magenta). Inserts show higher magnification of the dendrites highlighted with a white dashed line, which are presented in Figure 3B. Scale bar = 20 μm. **(C)** Full blot for GluA2/3 from the experiment presented in Figure 3C. Arrowheads indicate total and intracellular bands, arrows indicate crosslinked plasma membrane bands. **(D)** Full blot for GluA1 from the experiment presented in Figure 3C. Arrowheads indicate total and intracellular bands, arrows indicate crosslinked plasma membrane bands.

**Supplementary figure related to Figure 5:**
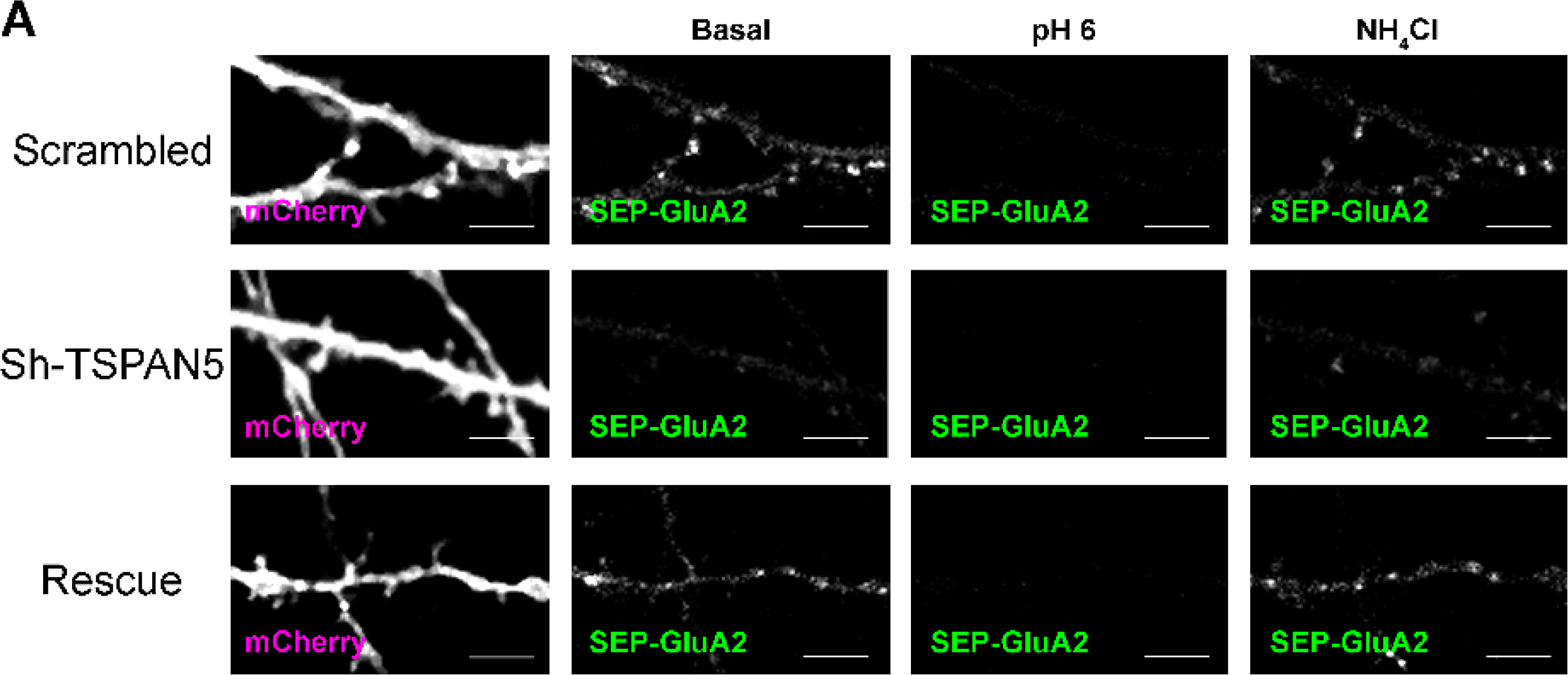
**(A)** Live confocal images of DIV20 culture rat hippocampal neurons transfected with SEP-GluA2 (green) and either Scrambled, Sh-TSPAN5 or Rescue construct co-expressing mcherry (magenta). Neurons were imaged in basal conditions, after exposure to pH 6 imaging media or after exposure to media containing 5 mM NH4Cl, to alkalinise intracellular compartments. Scale bar = 5 μm

## Notes

### Competing Interest Statement

The authors have declared no competing interest.

